# Characterization of a Human Liver NIST Reference Material fit for Proteomics Applications

**DOI:** 10.1101/834663

**Authors:** W. Clay Davis, Lisa E. Kilpatrick, Debra L. Ellisor, Benjamin A. Neely

## Abstract

The National Institute of Standards and Technology (NIST) is creating new, economical, qualitative reference materials and data for proteomics comparisons, benchmarking and harmonization. Here we describe a large dataset from shotgun proteomic analysis of RM 8461 Human Liver for Proteomics, a reference material being developed. Consensus identifications using multiple search engines and sample preparations demonstrate a homogeneous and fit-for-purpose material that can be incorporated into automated or manual sample preparation workflows, with the resulting data used to directly assess complete sample-to-data workflows and provide harmonization and benchmarking between laboratories and techniques. Data are available via PRIDE with identifier PXD013608.

## Background & Summary

NIST has a long history of developing reference materials (RMs)^1^ and advancing the measurement science used to characterize advanced materials. The advancement of non-targeted analysis (e.g. identification analysis, post-translational modification identification, differential analysis) and the need for RMs that are suitable for these types of measurements have fueled the generation of new materials and their associated data. With the rapid forward progress of instrumental technologies and continued innovations in bioinformatics, qualitative RMs and reference data sets directly linked to the RMs are a valuable resource for developing, improving, and assessing performance of methods, instrumental capabilities, and data processing tools.^2–4^

Linking data directly with a stable homogeneous material allows for the potential development of benchmarks of quality control metrics for peptide identification, bioinformatics, and systematic workflow evaluation. The development and availability of a common material for quality control purposes would significantly improve the critical evaluation of sample comparisons and processing and the variability of results.^5–7^

We report a data set of peptide identifications from RM 8461 Human Liver for Proteomics, a cryogenically homogenized and freeze-dried liver tissue developed as a material for complex proteomic analysis. The data set was collected using high resolution LC/MS instrumentation typically utilized in top-down and bottom-up protein analysis. The data set can serve as a resource for learning and training those new to the field as well as serve as a representative baseline proteomics data set for workflow analysis and bioinformatics. This data represents general laboratory preparation methods that can be used for evaluation of new or more complex methods (e.g. different enzyme digestions, depletion strategies, 1D or 2D LC). Additionally, since the processed data set used simple search parameters, the raw data can be re-analysed for additional peptide modifications and post-translational modifications.

There were three main concerns we sought to address in this analysis: homogeneity (reproducible protein inference via peptide identification), degradation of the proteome due to material preparation, and fit-for-purpose of the material for proteomic analysis. To address homogeneity, a stratified random sampling plan from across the RM production consisting of eight vials of RM 8461 were selected for analysis. Both 1 mg and 10 mg sub-sampling of the material from each of the eight vials was used, and the commonly identified peptides (and protein inference) evaluated using bottom-up proteomic analysis from a random stratified sample scheme across the RM production. Sampling and sample preparation at both the 1 mg and 10 mg sample size demonstrates the utility of the material for complex proteomic analysis. Semi-tryptic analysis with both Sequest HT and Mascot show relatively low sample decomposition and a high degree of protein integrity. Byonic-Preview also showed low levels of ragged N-term and C-term peptides as well as percentages that are similar to a commonly used commercial HeLa digest standard. Relative spectral abundance at both the peptide and protein level were highly correlated (e.g. reproducible) within all sample preparations utilized. The protein mass fraction, estimated by spectral abundance, yields detection and identification of peptides and protein inference of over several orders of magnitude within the sample. Despite the sample complexity, sample mass, or digestion method, there was a large number of commonly identified proteins (2055) between all of the Fusion Lumos runs. Comparison of the 1 mg sample runs with multiple search algorithms also showed a large number of commonly identified proteins (2703). The high level of commonly identified peptides and proteins and low level of proteome degradation demonstrate the fit-for-purpose of the material for proteomics analysis.

These results show that the RM is fit for purpose to perform QC when proteome coverage is being assessed. The material is complex enough to be used by other laboratories to assess performance of the sample preparation protocol, instrument performance, and database search parameters. Raw data will also prove valuable in development of specialized spectral libraries and software. As more data become publically available, users will be able to compare and assess protein measurements. This includes an improved ability to identify current best practices and challenges, ideally moving the fields toward harmonization.

## Methods

### Sample Material

A pool of liver material was cryogenically homogenized, freeze-dried, and radiation sterilized, limiting any proteolytic enzyme activity. The liver material was mixed, and aliquoted into approximately 0.5 g portions. Two sets of 8 vials were randomly selected across the production batch and tested for the new candidate RM 8461 Human Liver for Proteomics.

### Evaluation of Sample Preparation Methods

#### RapiGest/DTT/IAA

Approximately 1 mg (exact mass known) of candidate RM 8461 Human Liver for Proteomics was subsampled in triplicate from a single jar into 1.5 mL LoBind microcentrifuge tubes. The proteins were solubilized (pipet mixing) with an appropriate amount of 0.1 % mass fraction RapiGest (Waters, lot # 163011) in 50 mmol/L ammonium bicarbonate (Fluka) resulting in a 10 µg/µL solution. Next, 5 µL of each solution was added to 35 µL of 0.1 % mass fraction RapiGest and sonicated for 15 min, cooled on ice for 1 min, and then sonicated for another 15 min followed by the addition of 40 µL 50 mmol/L ammonium bicarbonate (Fluka). The samples were reduced with 10 µL of 45 mmol/L dithiothreitol (DTT; Sigma Aldrich; final concentration of 5 mmol/L) and incubated in an incubating shaker at 60 °C for 30 min then allowed to cool to room temperature. The mixture was alkylated using 3.75 µL of 375 mmol/L iodoacetamide (IAA, Pierce, Thermo Scientific; final concentration of 15 mmol/L) and incubated in the dark at room temperature for 30 min. Prior to trypsin addition, 100 µL of 50 mmol/L ammonium bicarbonate was added. A 1 µL aliquot of trypsin (Pierce Scientific, MS-Grade; 1 µg/µl in 50 mmol/L acetic acid; Fisher Scientific) was added (final mass ratio of 1:50 trypsin:protein) to each sample and incubated overnight at 37 °C. The digestion was halted and RapiGest cleaved with the addition of 100 µL 3 % volume fraction trifluoroacetic acid (TFA, Sigma Aldrich; 1 % final concentration) and incubated at 37 °C for 30 min before centrifugation and removal of the supernatant. Samples were desalted and concentrated using Pierce C18 spin columns (8 mg of C18 resin; Thermo Scientific) according to manufacturer’s instructions. Each sample was processed in duplicate yielding at maximum 60 µg peptides. These solutions were evaporated to dryness in a vacuum centrifuge and the samples were reconstituted in 50 µL 95 % water 5 % acetonitrile for analysis.

#### RapiGest/DTT/CAA

Approximately 1 mg (exact mass known) of RM 8461 Human Liver for Proteomics subsampled in triplicate from a single jar into 1.5 mL LoBind microcentrifuge tubes. The proteins were solubilized (pipet mixing) with an appropriate amount of 0.1 % mass fraction RapiGest, resulting in a 10 µg/µL solution. Next, 5 µL of each solution was added to 35 µL of 0.1 % mass fraction RapiGest and sonicated for 15 min, cooled on ice for 1 min, and then sonicated for another 15 min followed by the addition of 40 µL 50 mmol/L ammonium bicarbonate. The samples were reduced with 10 µL of 45 mmol/L DTT (final concentration of 5 mmol/L) and incubated in an incubating shaker at 60 °C for 30 min and then allowed to cool to room temperature. The mixture was alkylated using 10 µL of 400 mmol/L 2-chloroacetamide (CAA, Thermo Scientific; final concentration of 40 mmol/L) and incubated in the dark at room temperature for 30 min. Prior to addition of trypsin, 100 µL of 50 mmol/L ammonium bicarbonate was added. A 1 µL aliquot of trypsin (1 µg/µl in 50 mmol/L acetic acid) was added (1:50 trypsin:protein) to each sample and incubated overnight at 37 °C. The digestion was halted and RapiGest cleaved with the addition of 100 µL 3 % volume fraction TFA (1 % final concentration) and samples were incubated at 37 °C for 30 min before centrifugation and removal of the supernatant. Samples were processed using Pierce C18 spin columns to manufacturer’s instructions. Each sample was processed in duplicate yielding at maximum 60 µg peptides. These solutions were evaporated to dryness in a vacuum centrifuge and the samples were reconstituted in 50 µL 95 % water 5 % acetonitrile for analysis.

#### RapiGest/TCEP/CAA

Approximately 1 mg (exact mass known) of RM 8461 Human Liver for Proteomics subsampled in triplicate from a single jar into 1.5 mL LoBind microcentrifuge tubes. The proteins were solubilized (pipet mixing) with an appropriate amount of 0.1 % mass fraction RapiGest resulting in a 10 µg/µL solution. Next, 5 µL of each solution was added to 35 µL of 0.1 % mass fraction RapiGest and sonicated for 15 min, cooled on ice for 1 min, and then sonicated for another 15 min followed by the addition of 40 µL 50 mmol/L ammonium bicarbonate. The samples were reduced with 10 µL of 200 mmol/L Tris(2-carboxyethyl)phosphine hydrochloride (TCEP, Thermo Scientific; final concentration of 20 mM) and incubated at room temperature for 30 min. The mixture was alkylated using 10 µL of 400 mmol/L CAA (final concentration of 40 mmol/L) and incubated in the dark at room temperature for 30 min. Prior to addition of trypsin, 100 µL of 50 mmol/L ammonium bicarbonate was added. A 1 µL aliquot of trypsin (1 µg/µl in 50 mmol/L acetic acid) was added (1:50 trypsin:protein) to each sample and incubated overnight at 37 °C. The digestion was halted and RapiGest cleaved with the addition of 100 µL 3 % volume fraction TFA (1 % final concentration) and samples were incubated at 37 °C for 30 min before centrifugation and removal of the supernatant. Samples were processed using Pierce C18 spin columns according to manufacturer’s instructions. Each sample was processed in duplicate yielding at maximum 60 µg peptides. These solutions were evaporated to dryness in a vacuum centrifuge and the samples were reconstituted in 50 µL 95 % water 5 % acetonitrile for analysis.

#### SDC/TCEP/IAA

Approximately 1 mg (exact mass known) of RM 8461 Human Liver for Proteomics subsampled in triplicate from a single jar into 1.5 mL LoBind microcentrifuge tubes. The proteins were solubilized (pipet mixing) with an appropriate amount of 1 % mass fraction sodium deoxycholate (SDC; Thermo Scientific, lot # SJ2450944) in 50 mmol/L ammonium bicarbonate resulting in a 10 µg/µL solution. Next, 5 µL of each solution was added to 20 µL of 1 % mass fraction SDC in 50 mmol/L ammonium bicarbonate and sonicated for 15 min, cooled on ice for 1 min, and sonicated for another 15 min and then placed in an incubating shaker at 60 °C for 10 min. The samples were reduced with 2 µL of 200 mmol/L TCEP (final concentration of 14.8 mmol/L) and incubated at room temperature for 30 min. The mixture was then alkylated using 3.75 µL of 375 mmol/L IAA (final concentration of 37.5 mmol/L) and incubated in the dark at room temperature for 30 min. Prior to addition of trypsin, 170 µL of 50 mmol/L ammonium bicarbonate was added. A 1 µL aliquot of trypsin (1 µg/µl in 50 mmol/L acetic acid) was added (1:50 trypsin:protein) to each sample and incubated overnight at 37 °C. The digestion was halted with the addition of 100 µL 3 % volume fraction TFA (1 % final concentration). SDC removal was performed by four subsequent liquid-liquid extractions with 300 µL of ethyl acetate (Fisher Scientific), shaking by hand, centrifugation to promote phase separation, and removal of the upper ethyl acetate layer with the final removal of ethyl acetate performed with a speedvac. Samples were processed using Pierce C18 spin columns according to manufacturer’s instructions. Each sample was processed in duplicate yielding at maximum 60 µg of crude peptides. These solutions were evaporated to dryness in a vacuum centrifuge then the samples were reconstituted in 50 µL 95 % water 5 % acetonitrile for analysis.

#### SDC/TCEP/CAA

Approximately 1 mg (exact mass known) of RM 8461 Human Liver for Proteomics subsampled in triplicate from a single jar into 1.5 mL LoBind microcentrifuge tubes. The proteins were solubilized (pipet mixing) with an appropriate amount of 1 % mass fraction SDC in 50 mmol/L ammonium bicarbonate resulting in a 10 µg/µL solution. Next, 5 µL of each solution was added to 20 µL of 1 % mass fraction SDC in 50 mmol/L ammonium bicarbonate and sonicated for 15 min, cooled on ice for 1 min, and sonicated for another 15 min and then placed in an incubating shaker at 60 °C for 10 min. The samples were reduced with 2 µL of 200 mmol/L TCEP (final concentration of 14.8 mmol/L) and incubated at room temperature for 30 min. The mixture was then alkylated using 3 µL of 400 mmol/L CAA (final concentration of 40 mmol/L) and incubated in the dark at room temperature for 30 min. Prior to addition of trypsin, 170 µL of 50 mmol/L ammonium bicarbonate was added. A 1 µL aliquot of trypsin (1 µg/µl in 50 mmol/L acetic acid) was added (1:50 trypsin:protein) to each sample and incubated overnight at 37 °C. The digestion was halted with the addition of 100 µL 3 % volume fraction TFA acid (1 % final concentration). SDC removal was performed by four subsequent liquid-liquid extractions with 300 µL of ethyl acetate, shaking by hand, centrifugation to promote phase separation, and removal of the upper ethyl acetate layer with the final removal of ethyl acetate performed with a speedvac. Samples were processed using Pierce C18 spin columns according to manufacturer’s instructions. Each sample was processed in duplicate yielding at maximum 60 µg of crude peptides. These solutions were evaporated to dryness in a vacuum centrifuge then the samples were reconstituted in 50 µL 95 % water 5 % acetonitrile for analysis.

#### 10 mg Samples

A stratified random sampling plan of eight vials from across the candidate RM 8461 production run were selected for analysis. Approximately 10 mg (exact mass known) of RM 8461 Human Liver was subsampled from eight randomly selected vials of the material into individual 1.5 mL LoBind microcentrifuge tubes. The proteins were solubilized (pipet mixing) with an appropriate amount of 1 % mass fraction sodium deoxycholate (SDC; Thermo Scientific, lot # SJ2450944) in 50 mmol/L ammonium bicarbonate resulting in a 10 µg/µL solution. Next, 5 µL of each solution was added to 20 µL of 1 % mass fraction SDC in 50 mmol/L ammonium bicarbonate and sonicated for 15 min, cooled on ice for 1 min, sonicated for another 15 min and then incubated in an incubating shaker at 60 °C for 10 min. The samples were reduced with 2 µL of 200 mmol/L Tris(2-carboxyethyl)phosphine hydrochloride (TCEP, Thermo Scientific; final concentration of 14.8 mmol/L) and incubated at room temperature for 30 min. The mixture was then alkylated using 3 µL of 400 mmol/L 2-chloroacetamide (CAA, Thermo Scientific; final concentration of 40 mmol/L) and incubated in the dark at room temperature for 30 min. Prior to addition of trypsin, 170 µL of 50 mmol/L ammonium bicarbonate was added. A 1 µL aliquot of trypsin (1 µg/µl in 50 mM acetic acid) was added (1:50 trypsin:protein) to each sample and incubated overnight at 37 °C. The digestion was halted with the addition of 100 µL 3 % volume fraction trifluoroacetic acid (1 % final concentration). SDC removal was performed by four subsequent liquid-liquid extractions with 300 µL of ethyl acetate, shaking by hand, centrifugation to promote phase separation, and removal of the upper ethyl acetate layer with the final removal of ethyl acetate performed with a speedvac. Samples were processed using Pierce C18 spin columns, washed with 5 % volume fraction acetonitrile, and eluted with 70 % volume fraction acetonitrile. Each sample was processed in duplicate yielding at maximum 60 µg peptides. These solutions were evaporated to dryness in a vacuum centrifuge then the samples were reconstituted in 50 µL 95 % water 5 % acetonitrile for analysis.

#### 1 mg Samples

In order to evaluate the homogeneity on a smaller sample scale, triplicate preparations of 1 mg (exact mass known) of RM 8461 Human Liver was subsampled from the eight randomly selected vials of the material into individual 1.5 mL LoBind microcentrifuge tubes. The proteins were solubilized (pipet mixing) with an appropriate amount of 1 % mass fraction SDC in 50 mmol/L ammonium bicarbonate resulting in a 10 µg/µL solution and processed as the 10 mg samples described above.

#### NIST Gaithersburg Samples

An additional set of stratified random sampling plan of eight vials from across the candidate RM 8461 production run were selected for analysis to further assess the materials homogeneity. The digestion procedure was adapted from the protocol provided with Promega trypsin and is described below. Fifty mmol/L TCEP was prepared in 50 mmol/L Trizma buffer at pH 8.0 (adjusted with 5 mol/L NaOH). Trifluroethanol (TFE) was diluted to 56 % by volume in 50 mmol/L Trizma buffer, pH 8.3. One mg of liver was removed from each vial, weighed, and placed into low binding tubes (Eppendorf, Hauppauge, NY). Twenty µL of 50 mmol/L Trizma buffer (pH 8.3), 4.4 µL of 50 mmol/L TCEP and 20 µL of 56 % TFE were added. The samples were heat denatured at 60 °C for 60 min. Four µL of 300 mmol/L iodoacetamide was added and samples were vortexed and incubated in the dark for 45 min. Trypsin was prepared by adding 200 µL of Trizma buffer to 20 µg of trypsin. The 20 µg of trypsin was added to each sample. Digestions were performed at 37 °C for 20 hours. Digestions were stopped by adding 5 µL of a volumetric ratio of 50 % trifluoroacetic acid (final concentration was 0.5 % by volume). Final protein concentrations were estimated to be approximately 6 µg/µL.

#### High-pH Samples

Two full sets of the triplicate sample preparations were combined after 1D LC/MS/MS analysis and processed for 2D high-pH / reverse phase LC/MS/MS analysis. Briefly, the combined sample was analysed using a Dionex LC coupled to Foxy 200 fraction (Isco) collector. The peptide mixture (100 µL) was loaded onto a Zorbax 300 Extend C18 column (2.1mm id x 150 mm length, 3.5 µm particle size; Agilent Scientific) separated along a 70 min gradient of 100 % mobile phase A (5 % acetonitrile 10 mmol/L ammonium bicarbonate pH 8) and 0 % to 45 % mobile phase B (80 % acetonitrile 10 mmol/L ammonium bicarbonate pH 8) over 50 min, followed by a ramp to 100 % mobile phase B over 10 min at a flow rate of 250 µL/min with fractions collected every 90 s. Each fraction was then evaporated to dryness in a vacuum centrifuge and reconstituted in 20 µL 95 % water 5 % acetonitrile for LC/MS/MS analysis.

#### Instrumental Methods

For the LC/MS/MS analysis, the samples were analysed using an UltiMate 3000 Nano LC coupled to an Orbitrap Fusion Lumos mass spectrometer. Resulting peptide mixtures (1 µL) were loaded onto a PepMap 100 C18 trap column (75 µm id x 2 cm length; Thermo Fisher Scientific) at 3 µL/min for 10 min with 2 % acetonitrile and 0.05 % trifluoroacetic acid followed by separation on an Acclaim PepMap RSLC 2 µm C18 column (75µm id x 25 cm length; Thermo Fisher Scientific) at 40 °C. Peptides were separated along a 130 min gradient of 5 % to 27.5 % mobile phase B (80 % acetonitrile, 0.08 % formic acid) over 105 min followed by a ramp to 40 % mobile phase B over 15 min and lastly to 95 % mobile phase B over 10 min at a flow rate of 300 nL/min. The mass spectrometer was operated in positive polarity and data dependent mode (topN, 3 sec cycle time) with a dynamic exclusion of 60 sec (with 10 ppm error). The RF lens was set at 30 %. Full scan resolution was set at 120,000 and the mass range was set to m/z 375-1500. Full scan ion target value was 4.0 x 105 allowing a maximum injection time of 50 ms. Monoisotopic peak determination was used, specifying peptides and an intensity threshold of 1.0 x 10e4 was used for precursor selection. Data-dependent fragmentation was performed using higher-energy collisional dissociation (HCD) at a normalized collision energy of 32 with quadrupole isolation at m/z 0.7 width. The fragment scan resolution using the orbitrap was set at 30,000, m/z 110 as the first mass, ion target value of 2.0 x 105 and 60 ms maximum injection time.

#### Protein Search Parameters

Resulting raw files were processed and searched with Mascot (v2.6.2) for public upload and data archival on PRIDE^8^ via ProteomeXchange^9^. Raw files were additionally processed using all, or a combination of Sequest HT, Mascot, MSPepSearch, MSAmanda, and Byonic (ProteinMetrics; v3.2.0) algorithms via Proteome Discoverer (PD; v2.2.0.388).

Resulting raw files were processed and searched with PD using Sequest HT, Mascot, MSAmanda, Byonic, and MSPepSearch algorithms. Since some of the fractions following reverse phase high pH contained little to no peptides, only fractions 2 through 42 were used for searching. For Mascot searches, the UniProtKB SwissProt and SwissProt varsplic database (2018_06 release) was used and *Homo sapiens* was specified in the search parameters. For Sequest HT, MSAmanda, and Byonic searches, the *Homo sapiens* database (taxon ID: 9606) was retrieved from the 2018_06 release of the UniProtKB SwissProt database along with the SwissProt varsplic database. For Sequest HT, Mascot, MSAmanda, and Byonic searches, a cRAP database (common Repository of Adventitious Proteins, v 2012.01.01; The Global Proteome Machine) was used as well. For the MSPepSearch searches, three different spectral libraries were used: the 2014_05_29_human, a CID ion trap based library; human_hcd_selected_part1 and human_hcd_selected_part2, and an HCD library compiled in 2016.

The following search parameters were used with all algorithms with the exception of MSPepSearch: trypsin was specified as the enzyme allowing for two mis-cleavages (or semitryptic with up to 9 missed cleavages for the tissue decomposition analysis); carbamidomethyl (C) was fixed and acetylation (protein n-term), deamidation (NQ), pyro-Glu (n-term Q), and oxidation (M) were specified as variable modifications; 10 ppm precursor mass tolerance and 0.02 Da fragment ion tolerance. Within Sequest the peptide length was specified as a minimum of 6 and maximum of 144 amino acids. Resulting peptide spectral matches (PSMs) from both Mascot and Sequest searches were validated using the percolator algorithm, based on q-values at a 5 % false discovery rate (FDR). For MSPepSearch searches the following search parameters were used: 40 ppm precursor mass tolerance and 0.5 Da fragment ion tolerance, and a minimum match factor of 300. The spectral library-based results were further processed using a fixed value PSM validator node with a specified Δ Cn of 0.05.

### Code availability

Raw data was processed using all, or a combination of Sequest HT, Mascot (v2.6.2), MSPepSearch, MSAmanda, and Byonic (ProteinMetrics; v3.2.0) algorithms via PD (v2.2.0.388).

## Data Records

Each dataset was acquired with the goal of determining measureable and reproducible identifications in the candidate RM. All raw proteomics MS data files (in Thermo .raw file format) used for protein identification and relative quantification have been deposited to PRIDE^8^ via the ProteomeXchange Consortium^9^ and can be accessed with the dataset identifier PXD013608 (Table 1 – data files).

**Table 1.** Samples and Experimental Files in the Dataset. Raw data: raw and processed data are deposited on ProteomeXchange/PRIDE under Project DOI: 10.6019/PXD013608

| ID | MS Files<br>Proteomics | Raw Data<br>LC/MS |
| --- | --- | --- |
| 1 mg Sample Set | 1mg_1_A_1 - 1mg_1_A_3<br>1mg_1_B_1 - 1mg_1_B_3<br>1mg_1_C_1 - 1mg_1_C_3<br>1mg_2_A_1 - 1mg_2_A_3<br>1mg_2_B_1 - 1mg_2_B_3<br>1mg_2_C_1 - 1mg_2_C_3<br>1mg_3_A_1 - 1mg_3_A_3<br>1mg_3_B_1 - 1mg_3_B_3<br>1mg_3_C_1 - 1mg_3_C_3<br>1mg_4_A_1 - 1mg_4_A_3<br>1mg_4_C_1 - 1mg_4_C_3<br>1mg_5_A_1 - 1mg_5_A_3<br>1mg_5_B_1 - 1mg_5_B_3<br>1mg_5_C_1 - 1mg_5_C_3<br>1mg_6_A_1 - 1mg_6_A_3<br>1mg_6_B_1 - 1mg_6_B_3<br>1mg_6_C_1 - 1mg_6_C_3<br>1mg_7_A_1 - 1mg_7_A_3<br>1mg_7_B_1 - 1mg_7_B_3<br>1mg_7_C_1 - 1mg_7_C_3<br>1mg_8_A_1 - 1mg_8_A_3<br>1mg_8_B_1 - 1mg_8_B_3<br>1mg_8_C_1 - 1mg_8_C_3 | PDX013608 |
| 10 mg Sample Set | 10mg_1_A_1 - 10mg_1_A_3<br>10mg_2_A_1 - 10mg_2_A_3<br>10mg_3_A_1 - 10mg_3_A_3<br>10mg_4_A_1 - 10mg_4_A_3<br>10mg_5_A_1 - 10mg_5_A_3<br>10mg_6_A_1 - 10mg_6_A_3<br>10mg_7_A_1 - 10mg_7_A_3<br>10mg_8_A_1 - 10mg_8_A_3 | PDX013608 |
| Gaithersburg Sample Set | G_Liver_RM_9_A_1<br>G_Liver_RM_10_A_1<br>G_Liver_RM_11_A_1<br>G_Liver_RM_12_A_1<br>G_Liver_RM_13_A_1<br>G_Liver_RM_14_A_1<br>G_Liver_RM_15_A_1<br>G_Liver_RM_16_A_1 | PDX013608 |
| High pH Sample Set | Liver_RM_High_pH_01 -<br>Liver_RM_High_pH_48 | PDX013608 |
| Sample Preparation Comparison Set | Liver_RM_Rapigest_DTT_CAA_1 -<br>Liver_RM_Rapigest_DTT_CAA_3<br>Liver_RM_Rapigest_DTT_IAA_1 -<br>Liver_RM_Rapigest_DTT_IAA_3<br>Liver_RM_Rapigest_TCEP_CAA_1 -<br>Liver_RM_Rapigest_TCEP_CAA_3<br>Liver_RM_SDC_TCEP_CAA_1 -<br>Liver_RM_SDC_TCEP_CAA_3<br>Liver_RM_SDC_TCEP_IAA_1 -<br>Liver_RM_SDC_TCEP_IAA_3 | PDX013608 |

### Evaluation of RM 8461 Integrity

Semi-tryptic analysis of the 10 mg samples was performed with both the Sequest HT and Mascot search algorithms. The searches included up to eight missed cleavages in order to simulate and detect potential protein degradation. Trypsin cleaves peptides on the C-terminal side of lysine and arginine (if a proline is on the carboxyl side of the cleavage site, the cleavage will not occur) and the semi-tryptic search allows for the identification of degraded peptides that do not “end” with a lysine and arginine residue. For both Sequest HT and Mascot search results, a lower total number of PSMs, peptide groups, proteins, and protein groups were found in the semi-tryptic analysis than in the fully tryptic analysis (Table 2). Byonic-Preview was also used to quickly evaluate sample integrity and digestion completeness which are shown in Table 3 along with a standard HeLa digest (Pierce Scientific) standard used as a routine instrumental QC check. While the tryptic digestions of the RM are higher in the percentage of missed cleavages, both the ragged N-term and ragged C-term percentages are similar. Though biologically different materials, the comparison of the degree of ragged N-term and C-term demonstrates the materials are utility for the determination of sample integrity as well as software validation.

**Table 2.**
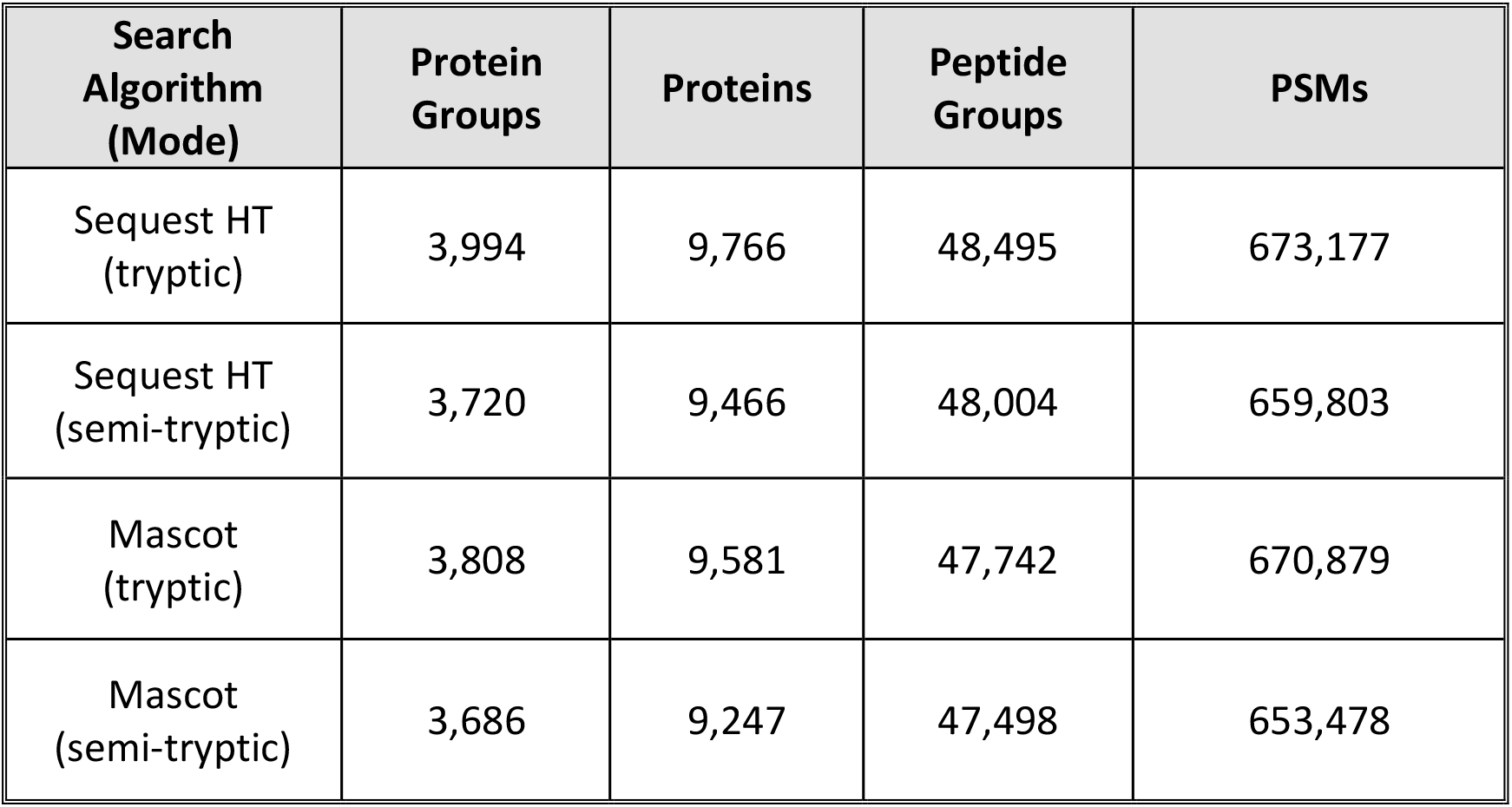
Comparison of Semi-tryptic and Tryptic Digestion Results with Sequest HT and Mascot Search Algorithms

| Search Algorithm (Mode) | Protein Groups | Proteins | Peptide Groups | PSMs |
| --- | --- | --- | --- | --- |
| Sequest HT (tryptic) | 3,994 | 9,766 | 48,495 | 673,177 |
| Sequest HT (semi-tryptic) | 3,720 | 9,466 | 48,004 | 659,803 |
| Mascot (tryptic) | 3,808 | 9,581 | 47,742 | 670,879 |
| Mascot (semi-tryptic) | 3,686 | 9,247 | 47,498 | 653,478 |

**Table 3.** Byonic-Preview Results of Trypsin Digestion from 10 mg Sample Preparations

| Sample | Missed Cleavage | Semi-tryptic Peptides ragged C-term | Semi-tryptic Peptides ragged N-term | Non-tryptic Peptides |
| --- | --- | --- | --- | --- |
| RM 8461 (n=8) | 15.0 % | 7.4 % | 1.1 % | 0.4 % |
| HeLa Digest Standard (n=4) | 5.8 % | 6.6 % | 2.4 % | 0.2 % |

### Reproducibility of Identified PSMs Utilizing Label-Free Quantification

The label free quantification node within PD 2.2 allows for relative peptide and protein quantification by integrating the extracted ion chromatogram from identified peptides (within 10 ppm mass error of peptide precursor *m/z*) and summing up individual peptide areas to a protein spectral abundance. Non-unique peptides belonging to multiple proteins are treated as such and their sums added to all resulting protein matches. This node allows for the qualification of not only the identified protein based on the precursor mass and fragment score to a primary sequence, but also a relative standard deviation (rsd) on replicate injections and sample preparations of the integrated peak areas determined by the extracted ion chromatograms. This allows for a filter of confidence on the reproducibility of instrument detection, and not just an algorithm derived identification. Triplicate injections of the 1 mg sample preparations show that 2374 of the 3682 commonly identified proteins have an integrated peak area from the resulting PSMs of less than 20 % rsd. In fact, the rsd across the triplicate injections from the triplicate sample preparations show 1136 of the commonly identified proteins with an integrated peak area from the PSMs of less of less than 20 % (Figure 1).

**Figure 1.**
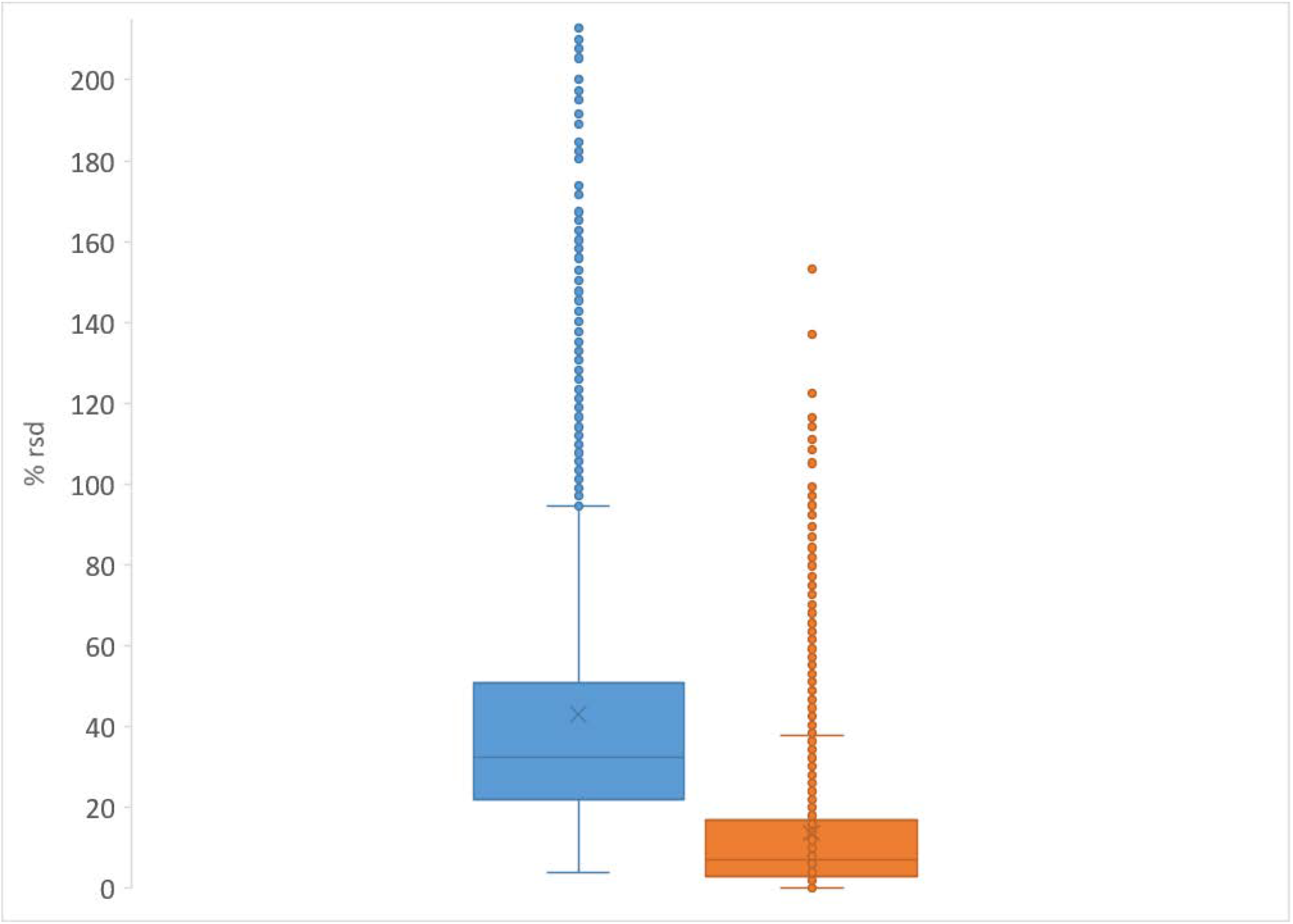
Range of % rsd from Commonly Identified Proteins (Blue – triplicate injections of triplicate sample preparation from 8 vials; Orange – triplicate injections from a single sample preparation from a single vial)

### Common Identifications – NIST Charleston and NIST Gaithersburg

Sample prepared from two sets of stratified random sampled vials across the production batch and replicate sample preparations identified 2055 common proteins using Sequest HT and analysed on the same instrument (Figure 2). While some of the individual 1D LC/MS results can yield between 3000 and 3500 protein identifications, the differing sample masses, sample preparation methods, and the number of vials sampled shows a significant correlation of identification homology and sample homogeneity. In addition, the protein intensity correlations between replicate sample preparations (Figure 3) is also well correlated and demonstrate relative quantitative similarities across samples.

**Figure 2.**
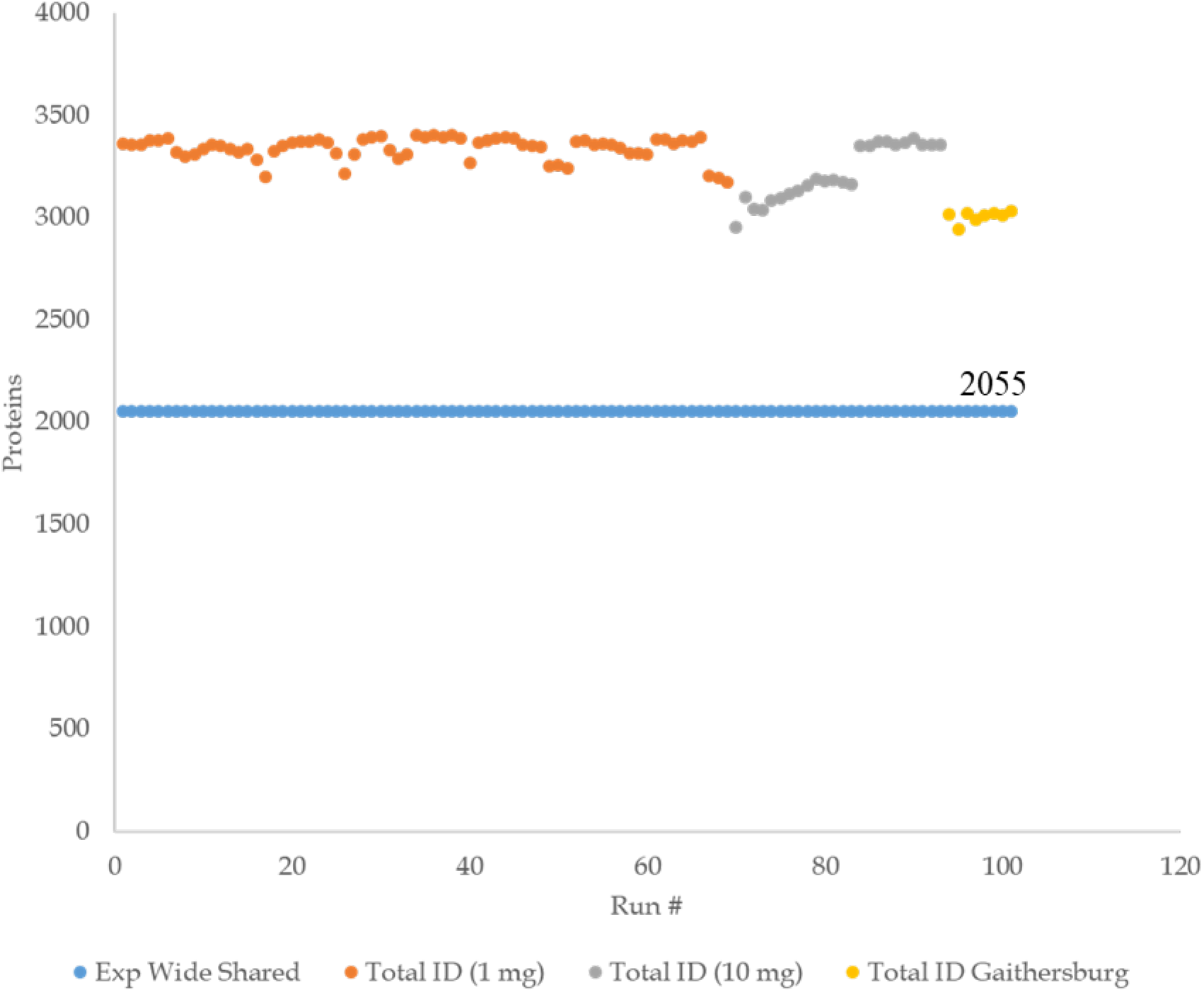
Total Number of Protein IDs and Experiment-Wide Shared Protein IDs

**Figure 3.**
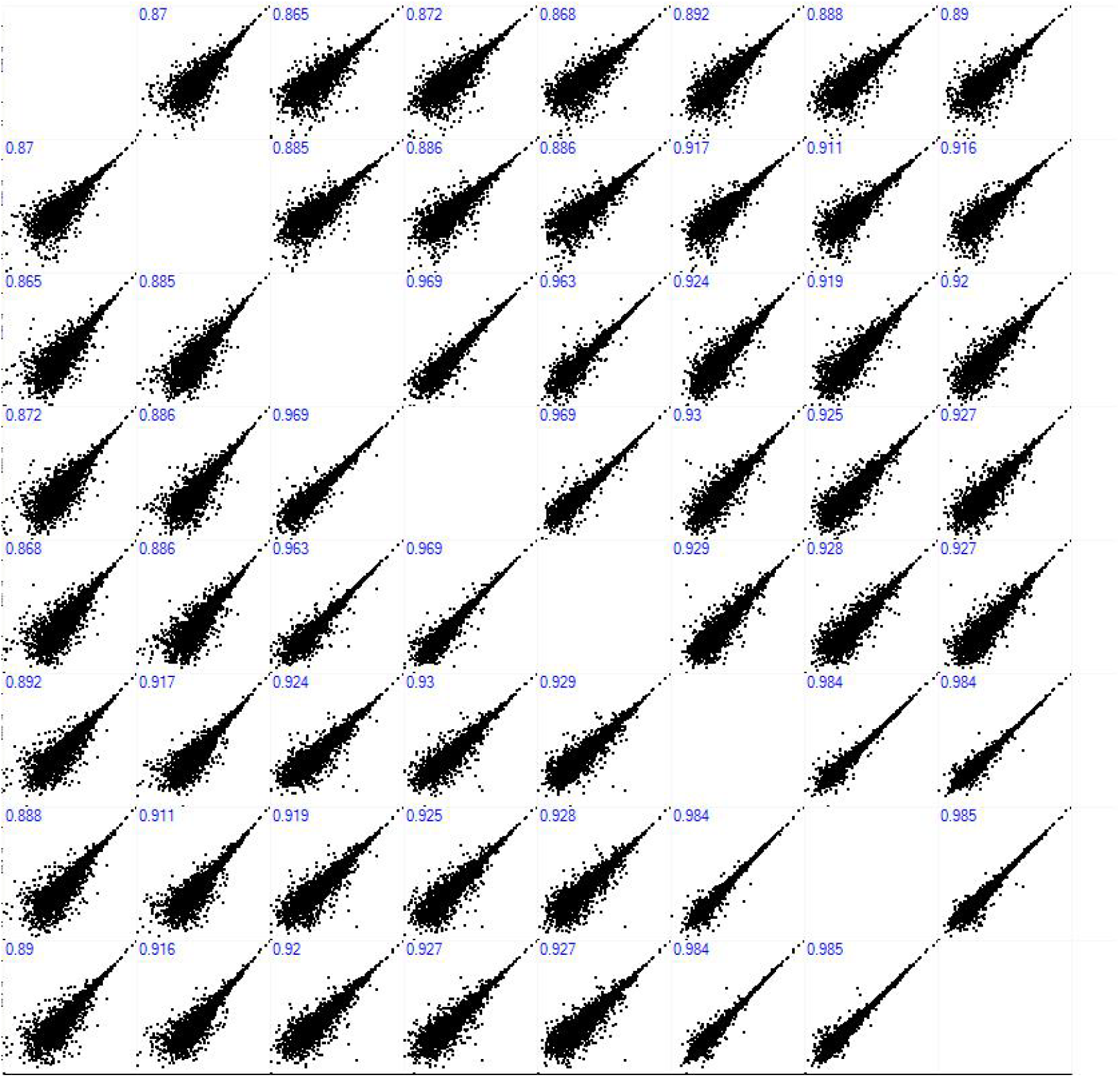
Protein Abundance Plots for Single Injections of Replicate Sample Preparations (10 mg) from Stratified Random Sampling of 8 Vials from RM 8461 Production

### Search Algorithm Comparison for Peptide and Protein Identification

Simultaneous and combined search results using the Sequest HT, Mascot, MSAmanda, MSPepSearch, and Byonic algorithms produced at total of 9,153,280 PSMs, 553,908 peptide groups, 19,778 proteins, and 7,873 protein groups resulting from over 591 hours of search time. The overlap of identified peptides (Figure 4) and proteins (Figure 5) among all of the search algorithms is quite significant. Upwards of 25,000 peptides are commonly identified resulting in almost 3000 common proteins. This number is reduced to 2693 proteins when an additional constraint of a minimum of 3 unique peptides are required for identification.

**Figure 4.**
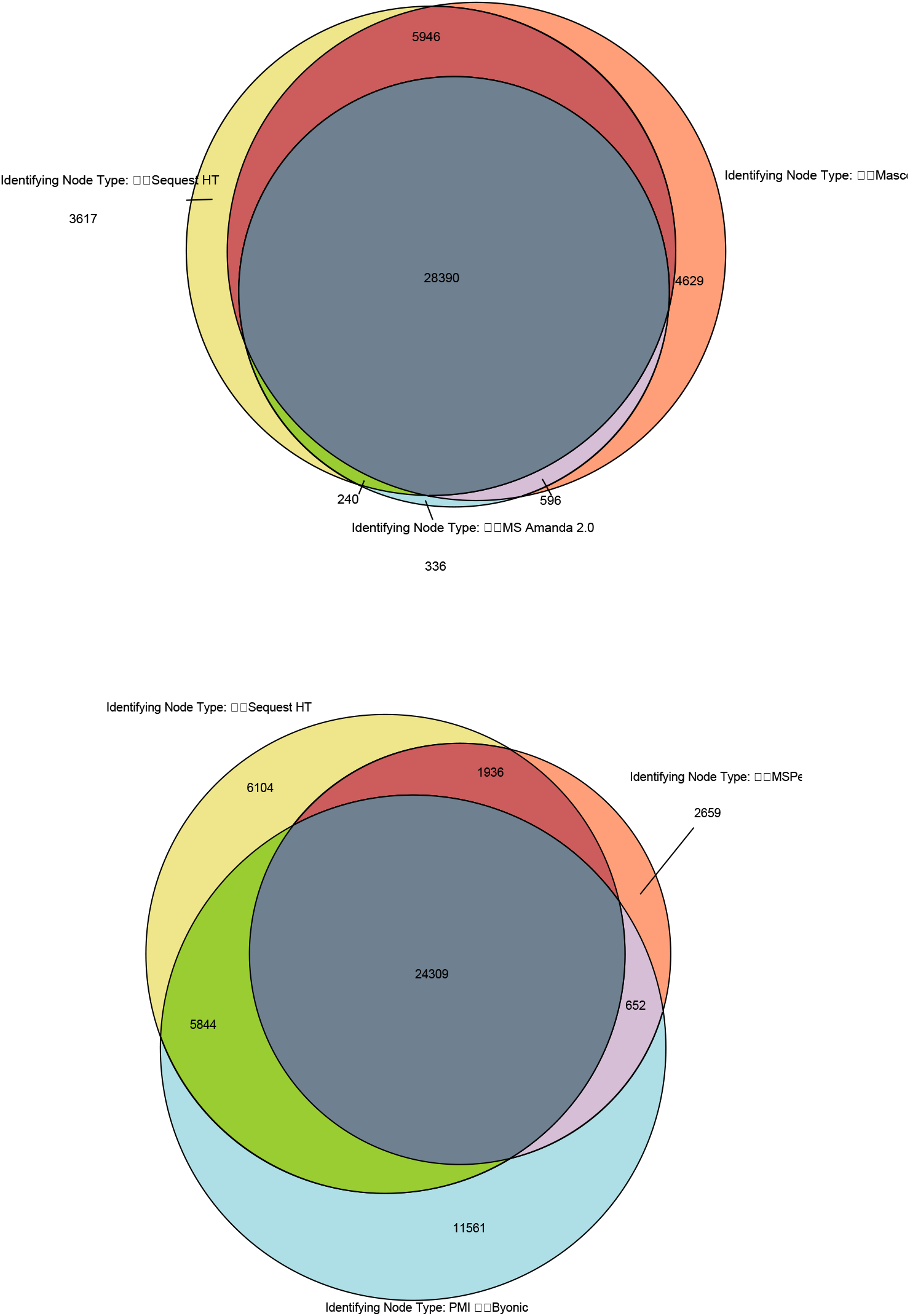
Venn Diagram of Commonly Identified Peptides across Search Algorithms

**Figure 5.**
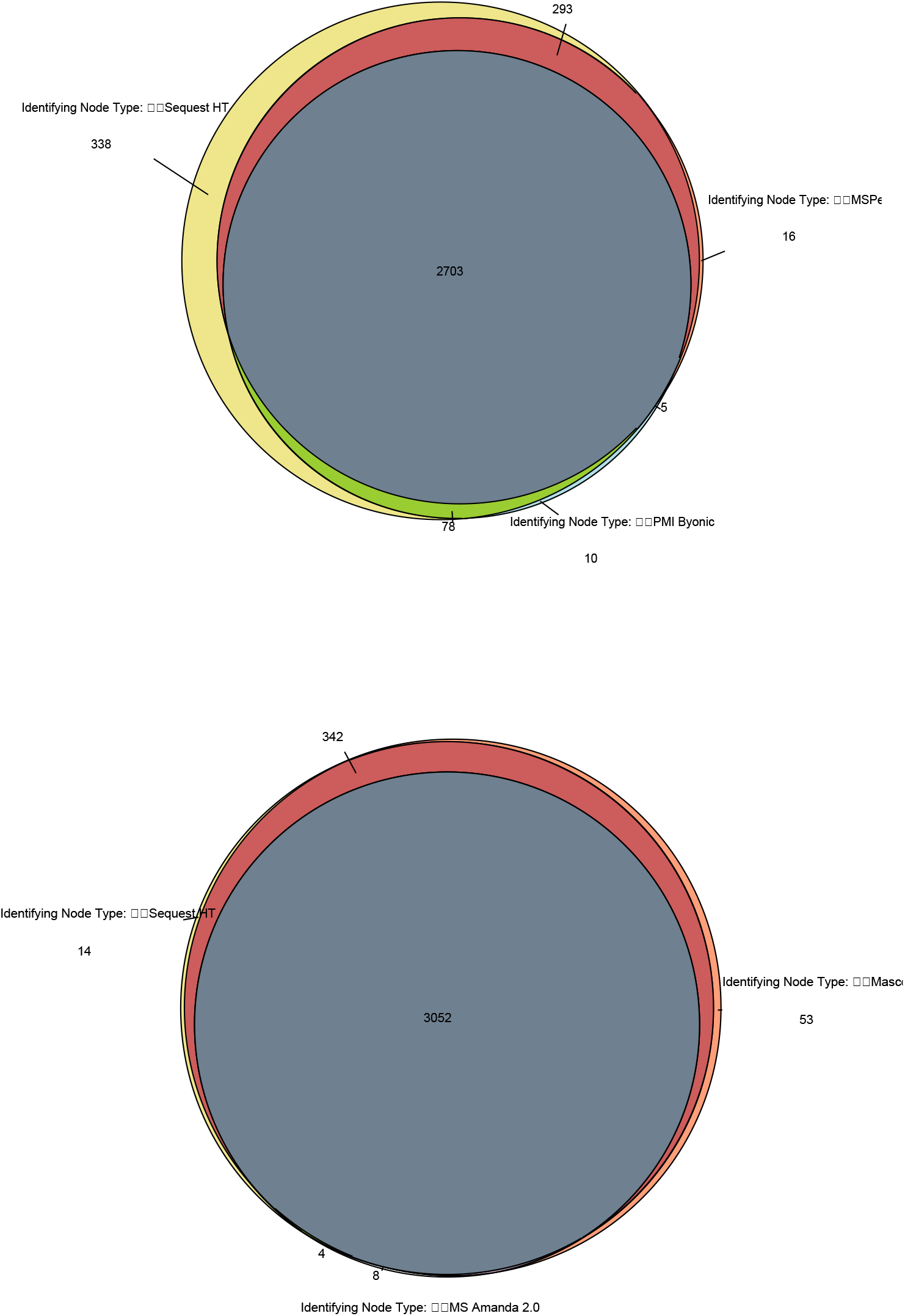
Venn Diagram of Commonly Identified Proteins across Search Algorithms

MSPepSeach results tend to give fewer identifications compared to standard search algorithm results due to the inherent difference in what is being searched and matched. MS/MS library identification matches are based on pairing an experimental peptide MS/MS with a MS/MS in the library and any differences between the reference and experimental spectra allowing for very rapid generation of search results. However, the results are restricted to annotated spectra in the reference library, which severely limits the identification of modified peptides (static and/or post-translational modifications).

### High pH 2D LC/MS/MS

In order to achieve a more comprehensive identification of proteomes, two-dimensional (2D) chromatography has been an invaluable tool.^10–12^ The 2D method demonstrate feasibility of HpH workflow using a small sample masses yielding high proteome coverages. Utilizing the 2D sample processing workflow show increases in all aspects of analysis resulting in the identification of 109,342 peptide groups corresponding to 21,847 proteins from 8407 protein groups with 1,047,269 PSMs. Figure 6 shows the abundance range of identified proteins over 7 orders of magnitude from the 2D analysis.

**Figure 6.**
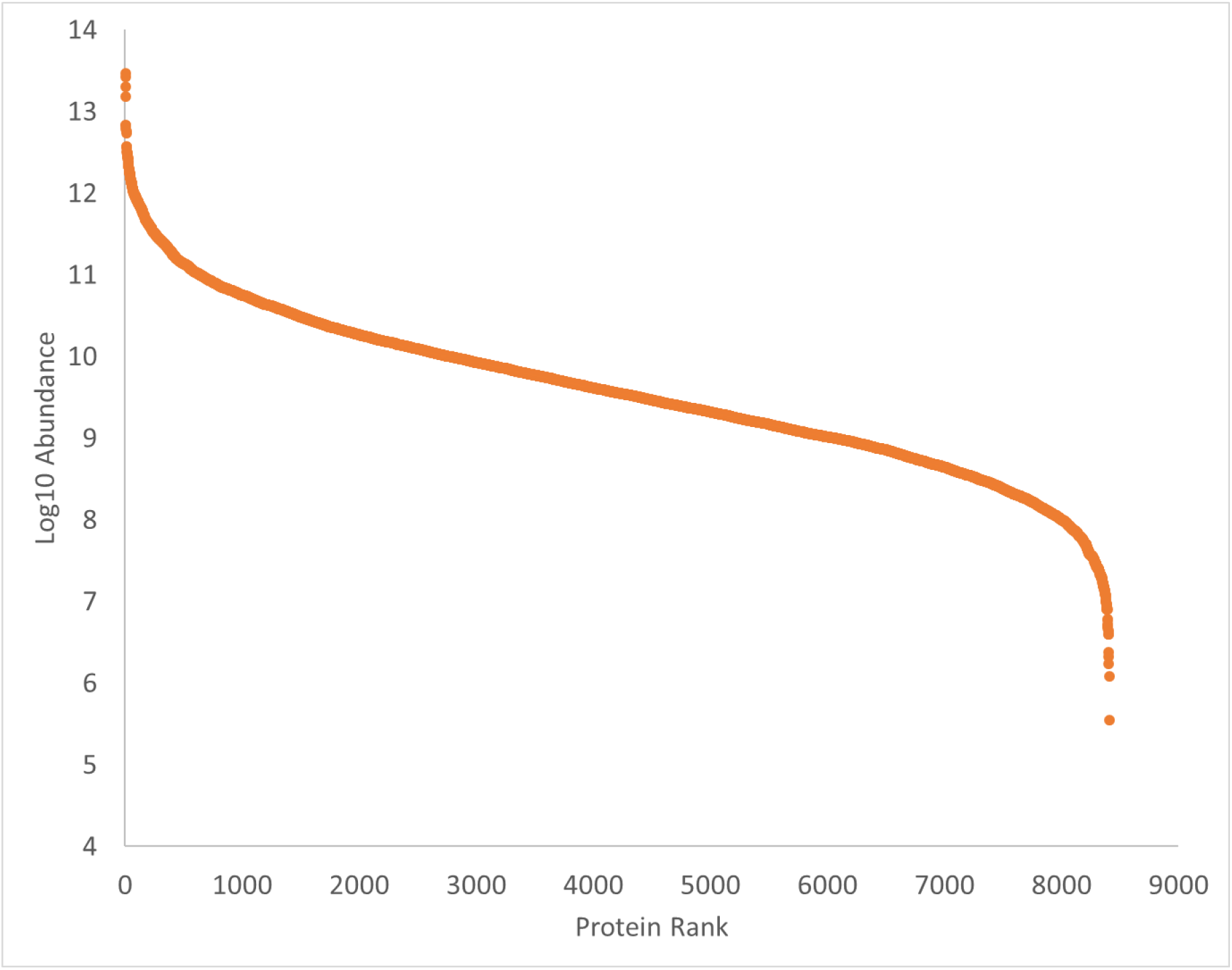
Protein Abundance Ladder for High pH 2D LC/MS/MS

## Technical Validation

Development of the liver material was intended to serve as QC material for bottom-up proteomic measurements utilizing standard processes and instrumentation. Identified proteins were selected at 5 % FDR, identified PSMs with reproducibility of less than 20 % rsd in replicate sample preparations, and multiple search algorithms to ensure data quality and material robustness. Instrument and LC separation parameters as well as data validation were checked for system suitability using HeLa standard protein digest standard. Homogeneity, in terms of reproducible protein inference via peptide identification, was assessed at 10 mg and 1 mg sample masses across the production batch using random stratified samples and shows a high degree of commutability as well as between sample preparation methods. The sample integrity of the materials was assessed and validated by semi-tryptic analysis with multiple search algorithms and is comparable to the commercially available HeLa digest.

Despite a much higher total number of identifications utilizing the high pH method (21,847 proteins), an overly stringent identification criteria was used to finalize a list of proteins whose prevalence and identification are assured in the sample regardless of approaches/platforms. First, commonly identified proteins from the 1 mg triplicate preparations from box 31-jar 29 and replicate injections with a spectral abundance reproducibility of less than 20 % CV were considered (1891). Additionally, only proteins identified with Sequest HT, Mascot, MSAmanda, and Byonic algorithms and having a minimum of 3 unique peptides were considered. When combined, a total of 1619 proteins met all of the above criteria (Table 3).

## Usage Notes

Though the main goal of the project was to develop qualitative reference materials and data for proteomics comparisons, benchmarking and harmonization and demonstrating fit-for-purpose of the material and data to serve as a benchmark for laboratories and users across platforms and methods. The data set can be re-searched with any bottom up bio-informatics platform, workflow, and with additional modifications (e.g., phosphorylation, glycosylation, ubiquitination, etc.) to glean more information on both the sample and data.

## Acknowledgements

The authors would like to acknowledge researchers from the NIST Environmental Specimen Bank Group, including Rebecca Pugh, Amanda Moors, and Jennifer Ness for their assistance in processing and bottling the liver material.

## Author contributions

W.C.D. designed experiments, performed LC/MS proteomics sample processing, data acquisition, data analysis, and wrote the manuscript. L.E.K designed experiments, performed LC/MS proteomics sample processing, data acquisition, data analysis, and wrote the manuscript. D.L.E. was responsible for pre-analytical work on the RM material, designed experiments, and wrote the manuscript. B.A.N. designed experiments, performed LC/MS proteomics data analysis, and wrote the manuscript.

## Competing interests

The authors declare no competing financial interests.

## Disclaimer

Official contribution of the National Institute of Standards and Technology. Not subject to copyright in the United States. Certain commercial equipment, instruments, software or materials are identified in this document. Such identification does not imply recommendation or endorsement by the National Institute of Standards and Technology, nor does it imply that the products identified are necessarily the best available for the purpose.

## Notes

https://www.ebi.ac.uk/pride/archive/projects/PXD013608

## References

1 Standard Sample No. 1, Argillaceous Limestone (United States Department of Commerce, Bureau of Standards, 1910).

2 Bittremieux, W. et al. Quality control in mass spectrometry-based proteomics. Mass Spectrometry Reviews 37, 697–711, doi:10.1002/mas.21544 (2018).

3 Sempos, C. T. et al. Vitamin D status as an international issue: National surveys and the problem of standardization. Scandinavian Journal of Clinical & Laboratory Investigation 72, 32–40, doi:10.3109/00365513.2012.681935 (2012).

4 Beasley-Green, A., Bunk, D., Rudnick, P., Kilpatrick, L. & Phinney, K. A proteomics performance standard to support measurement quality in proteomics. Proteomics 12, 923–931, doi:10.1002/pmic.201100522 (2012).

5 Ludwig, K. R., Schroll, M. M. & Hummon, A. B. Comparison of In-Solution, FASP, and S-Trap Based Digestion Methods for Bottom-Up Proteomic Studies. Journal of Proteome Research 17, 2480–2490, doi:10.1021/acs.jproteome.8b00235 (2018).

6 Glatter, T., Ahrne, E. & Schmidt, A. Comparison of Different Sample Preparation Protocols Reveals Lysis Buffer-Specific Extraction Biases in Gram-Negative Bacteria and Human Cells. Journal of Proteome Research 14, 4472–4485, doi:10.1021/acs.jproteome.5b00654 (2015).

7 Piehowski, P. D. et al. Sources of Technical Variability in Quantitative LC-MS Proteomics: Human Brain Tissue Sample Analysis. Journal of Proteome Research 12, 2128–2137, doi:10.1021/pr301146m (2013).

8 Vizcaino, J. A. et al. 2016 update of the PRIDE database and its related tools. Nucleic Acids Research 44, D447–D456, doi:10.1093/nar/gkv1145 (2016).

9 Vizcaino, J. A. et al. ProteomeXchange provides globally coordinated proteomics data submission and dissemination. Nature Biotechnology 32, 223–226, doi:10.1038/nbt.2839 (2014).

10 Dwivedi, R. C. et al. Practical implementation of 2D HPLC scheme with accurate peptide retention prediction in both dimensions for high-throughput bottom-up proteomics. Analytical Chemistry 80, 7036–7042, doi:10.1021/ac800984n (2008).

11 Bekker-Jensen, D. B. et al. An Optimized Shotgun Strategy for the Rapid Generation of Comprehensive Human Proteomes. Cell Systems 4, 587–+, doi:10.1016/j.cels.2017.05.009 (2017).

12 Kulak, N. A., Geyer, P. E. & Mann, M. Loss-less Nano-fractionator for High Sensitivity, High Coverage Proteomics. Molecular & Cellular Proteomics 16, 694–705, doi:10.1074/mcp.O116.065136 (2017).

